# Diagnostic of head blight and leaf blotch diseases of wheat in Sweden using a mycobiome isolation approach

**DOI:** 10.1101/2024.01.09.574850

**Authors:** Harleen Kaur, Elisa Vilvert, Miguel Ángel Corrales Gutiérrez, Jiasui Zhan, Dimitar Douchkov, Francesca Desiderio, Heriberto Vélëz, Salim Bourras

## Abstract

Increasing the durability of crop resistance to plant pathogens has become a major concern in plant pathology. There are 32 diseases of agronomical importance in wheat, which makes disease management in this crop of worldwide importance extremely challenging. Two important groups of wheat pathogens driving the consumption of chemical fungicides in Sweden and worldwide are those causing head blight diseases and leaf blotch diseases. Swedish fields are regularly infested with head blight diseases typically driven by a complex of *Fusarium* and non-fusarium taxa including *F. graminearum*, *F. culmorum*, while leaf blotches are typically caused by *Zymoseptoria tritici*, *Pyrenophora tritici-repentis*, and *Parastagonospora nodorum*. In this work, we present a survey of the larger diversity of taxa associated with head blight and leaf blotch diseases in Sweden, with the primary aim to assess the usefulness of such an approach to develop new understanding and tools for disease diagnostics and management *sensu lato*. Here, we analyze nearly a thousand isolates collected from wheat fields across eight counties in Sweden and report the identification of pathogenic and non-pathogenic taxa as well as candidate biological control agents associated with head blight and leaf blotch diseases. We argue that such surveys represent an important tool for improving disease diagnostics, disease forecasting, and integrated management in wheat.

## Introduction

Disease identification in the field can be very challenging, considering the complexity and the convergence of symptoms caused by different pathogen species. For diseases typically caused by a complex such as Fusarium Head Blight (FHB), several fungal species can be implicated. For instance, there are 17 *Fusarium* sp. and non-*Fusarium* species implicated in FHB. Of these, *F. graminearum* and *F. culmorum* are often presented as the most important drivers of the disease, with F*. graminearum* often being the most predominant species (**Karlsson et al., 2021**). The composition of the complex can significantly vary among countries depending on the varieties grown in the field, agricultural practices, or climate (**Karlsson et al., 2021**). For instance, a field monitoring study conducted by Xu et al. (2005) in four European countries (Hungary, Ireland, Italy, and the UK), it was shown that FHB was frequently associated with *F. graminearum* and *F. poae* in Italy and Hungary but not in Ireland and the UK, while *F. culmorum* was rarely detected except in Ireland (**Xu et al., 2005**). This demonstrates the importance of regularly monitoring FHB for the correct identification of the causal agents and proper assessing the efficiency of management strategies.

Other prominent examples of phenotypic complexity are leaf blotch diseases caused by *Zymoseptoria tritici*, *Pyrenophora tritici-repentis*, and *Parastagonospora nodorum*. All pathogens cause convergent disease symptoms that can be confounded with each other. Actually, the three pathogen species were historically confounded as one species, then *Z. tritici* and *P. nodorum* were referred to as the Septoria blotch complex or Septoria complex due to frequent co-occurrence of the two species and the similarity of symptoms. From a resistance breeding perspective, the distinction between those three species is crucial since it was shown that the host resistance is genetically different (i.e., requires different resistance genes, resistance mechanisms, and breeding strategies) (**Brown et al., 2015**). This further demonstrates the importance of monitoring and precise disease diagnostics as a tool for disease management. Furthermore, leaf blotch diseases in wheat are still largely controlled with fungicides, leading to the rapid development of fungicide resistance in pathogen populations across Europe (**Pereira et al., 2020; Justesen et al., 2021; Jørgensen et al., 2021, Jørgensen et al., 2022**) Thus, surveys are essential for the management of fungicide resistance and further highlight the importance of continuous isolation and characterization of pathogen populations from the field.

One major challenge for disease diagnosis in wheat is the recurrent emergence of hypervirulent pathogen races, with one prominent example being the stem UG99 from the wheat pathogen *Puccinia graminis* f. sp. *tritici*, that was responsible for a global pandemic of stem rust in wheat (**Singh et al., 2011**). From a phenotypic perspective, it is impossible to distinguish the stem rust symptoms caused by the UG99 race from any other race, thus highlighting the importance of molecular race characterization for proper monitoring and resistance breeding (**Patpour et al., 2022**). A similar challenge is imposed by *Blumeria graminis* f.sp. *triticale* (*B.g. triticale*), a new wheat pathogen that emerged from the hybridization of wheat powdery mildew (*B.g. tritici*) and rye powdery mildew (*B.g. secalis*) (**Menardo et al., 2016**). *B.g. triticale* is a new pathogen with an expanded host range that can not only infect the Triticale crop but also wheat, causing symptoms that are indistinguishable from the native wheat powdery mildew species *B.g. tritici* (**Menardo et al., 2016**) These examples further demonstrate the importance of monitoring and the molecular identification of the causal agent as a tool for the surveillance of emerging and re-emerging diseases.

The wheat mycobiome has been shown to be an important source of beneficial fungal endophytes with promising biological control properties. Such antagonists often occur in the same ecological niches/tissues as their ‘target’ pathogen species and have thus been isolated from various organs, including roots, leaves, seeds, or ears (**Huang & Pang, 2017; Yue et al., 2018; Rojas et al., 2020**). Considering the genetic diversity of the endophytic taxa inhabiting wheat, only a fraction of the wheat mycobiome has been characterized for its biological control properties. Such work is mainly restricted by the tedious initial isolation and molecular characterization of hundreds (if not thousands) of fungal taxa. We, therefore, argue that surveys for diseases could be coupled with surveys for biological control agents, taking advantage of the availability of host tissues where both pathogens and potential antagonists can be isolated. In this study, we aimed to provide an experimental proof-of-concept for the possible uses of disease sampling and monitoring campaigns for constructing a wheat mycobiome library that can be used to understand and monitor wheat head blight and leaf blotch diseases in Sweden. In particular, we assess if such a library can inform on shifts in pathogen populations and possibly lead to the identification of potential emerging diseases and candidate biological control agents.

## Materials and methods

### Sample collection and handling

A survey of wheat diseases was conducted in collaboration with the plant protection extension services of the Swedish Board of Agriculture. In 2020, spikes were collected from spring wheat varieties cultivated in Sweden and grown in a small propagation plot north of Stockholm. The plot consisted of the six Spring wheat varieties: Alderon, Dacke, Dala, Happy, Rohan, Stanley; two spring wheat genotypes used as scientific standards: Bobwhite and Fielder; and one triticale genotype. In 2021 and 2023, spike and leaf samples were collected by field pathologists from the plant protection extension services from eight counties across Sweden: Norrbotten, Uppsala, Stockholm, Södermanlands, Östergötland, Västra Götaland, Gottland, and Skåna (Figure 1 A). Samples were typically shipped within 24 hours in glycine bags and immediately processed or stored at 4°C in dry storage until further processing.

**Figure 1.**
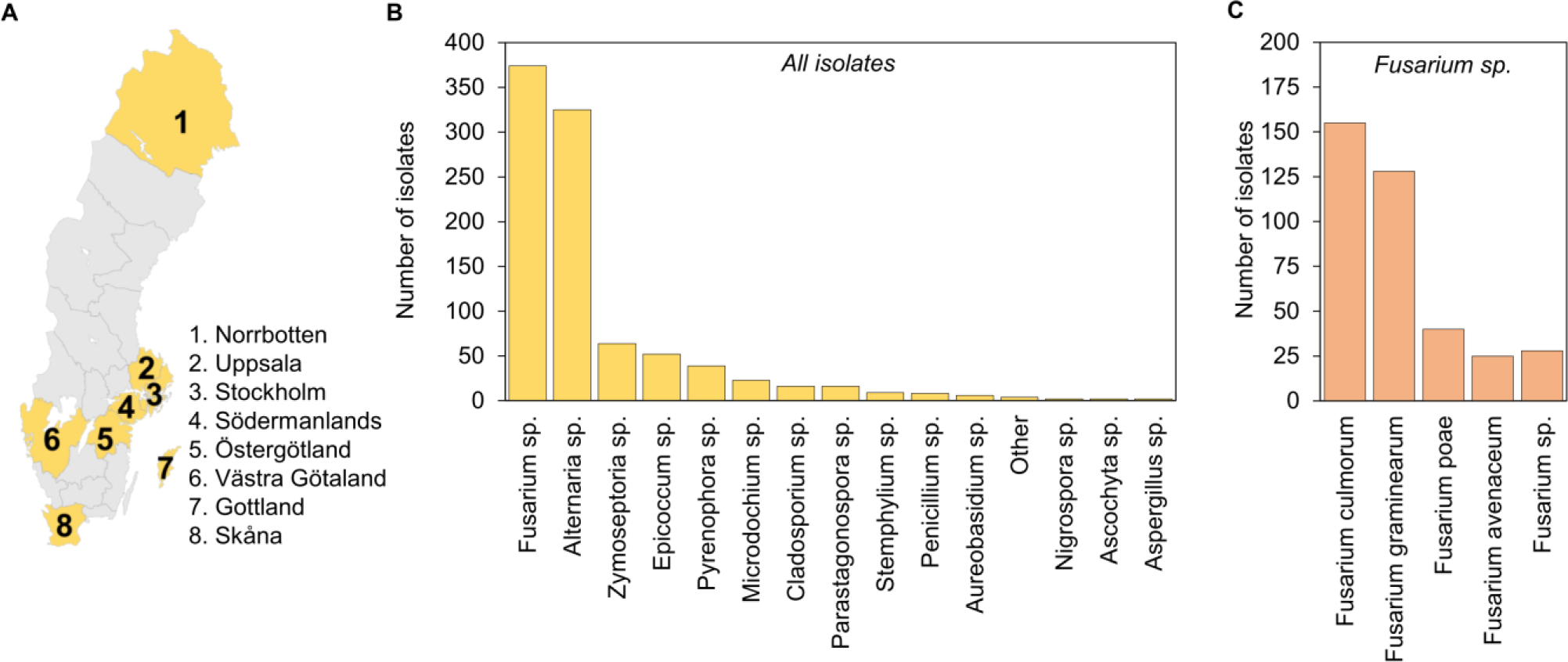
Summary and overall diversity of the wheat mycobiome library. (A) Geographical origin of the samples. (B) Overall diversity of the fungal genera represented in the library. (C) Partial analysis of species diversity of *Fusarium* sp.

### Media preparation for isolation and long-term conservation

Two different media were used for pathogen isolation, purification, and long-term storage. Potato dextrose agar (PDA) was prepared from dehydrated pre-mixed media from MP Biomedicals (USA) according to the manufacturer’s instructions. Yeast malt sucrose (YMS) agar medium was prepared by mixing solve 4 g yeast extract, 4 g malt extract, and 4 g sucrose, and 16 g agar in 1000 ml milliQ water, then autoclaving for 5 min at 121°C. For mid-term storage, isolates were kept as slow-growing cultures on half-strength PDA or YMS agar in small glass vials and stored in the dark at 4°C. For long-term storage, three to four ca, 5mm mycelial plugs were cut from a growing colon, placed in a 2ml cryotube containing 30% glycerol, and stored at −80°C. For *Z. tritici*, long-term storage vials were prepared from liquid cultures as previously described by Fagundes and collaborators (**Fagundes et al., 2020**).

### Isolation and purification of fungal colonies from spikes and leaves

Seeds and glumes were manually separated from full spikes and surface sterilized under a sterile hood for one minute in 1% sodium hypochlorite, followed by 30 seconds in 70% ethanol, and washed twice for 30 seconds in autoclaved Milli-Q water. Individual seeds were placed in Petri plates containing fungal culture media amended with 50 µg/ml of Kanamycin. For leaf segments a square of ca 5mm width was dissected around blotch symptoms typically consisting of necrotic spot often surrounded by a chlorotic ring. Segments were surface sterilized under a sterile hood for 30 seconds in sodium hypochlorite, followed by 30 seconds in 70% ethanol, and washed twice for 30 seconds in autoclaved Milli-Q water. Three samples from the same leaf were placed in Petri plates containing fungal culture media amended with 50 µg/ml of Kanamycin. Alternatively, samples were also simply washed in Milli-Q water and further processed into plating. All plates were incubated at 20℃ until colonies were visible (usually around 5 to 7 days). Individual colonies were picked and transferred to new plates and incubated at 20℃. The process was repeated until a single pure colony was obtained. For the specific case of *Z. tritici*, leaf segments harboring pycnidia were incubated at 18°C in a humid chamber consisting of a Petri dish supplemented with a sterile absorbent paper disk soaked with sterile water. Plates were inspected every day under a binocular to search for bursting pycnidia where mucilage had been ejected. In such a case, the pycnidium was carefully fetched using a sterile needle under a binocular placed inside a sterile hood and transferred into a Petri dish with YMS agar, supplemented with 50 µg/ml of Kanamycin.

### DNA extraction and genotyping by fragment sequencing

A modified cetyltrimethylammonium bromide (CTAB) based DNA extraction method was used to extract DNA of fungal isolates. Briefly, mycelium plugs were used to inoculate a liquid medium containing malt extracts and vitamins (MEV). Cultures were incubated with ca. 150rpm shaking in the dark at 25℃ for two weeks. Then, mycelium was harvested, homogenized with a CTAB buffer (3% CTAB; 150 mM TRIS-HCl; 2.6 M NaCl; 20 mM EDTA), and incubated at 65℃ for 30 min inside a water bath. The samples were centrifuged at 10’000 g for 10 min and the supernatant was transferred to a new tube supplemented with an equal volume of chloroform. After a second centrifugation at 10’000 g for 10 min, the supernatant was transferred to a new tube, an equal amount of isopropanol was added, and the mix was incubated at −20℃ for a few minutes. A final centrifugation at 10’000 g for 10 min resulted in the deposition of a DNA pellet, which was washed using 70% ethanol and resuspended in TE buffer. In addition, another DNA extraction method was used to extract the DNA of the fungal isolates. First, mycelium plugs from the isolates were cultured in falcon tubes containing 10 ml of YMS liquid medium and incubated for 7-15 days at 150 rpm and 25 °C in dark. Then, the mycelium was harvested, freeze-dried, and subjected to genomic DNA extraction using the Nucleo Mag Plant kit (Macherey-Nagel GmbH &Co. KG, Germany) in the automated nucleic acid extractor Maelstrom 4800 (TANBead Taiwan Advanced Nanotech Inc., Taiwan).

Identification of pathogen species was carried out by Polymerase Chain Reaction (PCR) of the genomic DNA followed by amplicon sequencing of conserved nuclear locus, internal transcribed spacer (ITS) rDNA. For ITS PCR, primer pair ITS1 (5’-TCCGTAGGTGAACCTGCGG-3’) and ITS4 (5-TCCTCCGCTTATTGATATGC-3’) were used for genotyping (**White T.J., 1990**). PCR amplification was followed by PCR clean-up using exonuclease kit (Thermo Fisher Scientific) according to the manufacturer’s instructions. Amplicons were subsequently sent for fragment sequencing at Macrogen Europe (The Netherlands, https://www.macrogen-europe.com/). DNA sequences were manually curated and analyzed for sequence homology to fungal sequences deposited at the National Center for Biotechnology Information (NCBI) using the BLASTn search tool. Isolates used for leaf infection assays were also tested with *F. graminearum* marker GOFW (5’-ACCTCTGTTGTTCTTCCAGACGG-3’) GORV (5’-CTGGTCAGTATTAACCGTGTGTG-3’) (de Biazio et al., 2008), and for *F. culmorum* with Fc03 (5’-TTCTTGCTAGGGTTGAGGATG-3’) Fc02 (5’-GACCTTGACTTTGAGCTTCTTG-3’) (**Astrid Bauer and Seigner, 2015**).

### Plant cultivation and infection assays

Wheat plants were grown in the phytotron (controlled climate chambers) for three to four weeks at 20℃, 70% relative humidity and 300µMOL light intensity. The inoculum was composed of isolate HK48 (*Fusarium graminearum*), HK50 (*Fusarium culmorum*). The isolates were incubated for two weeks at 25℃ and constant darkness on OMA. The conidia were scraped from the colony surface and resuspended in sterile water. The inoculum cell density was adjusted to 250 spores per µl using water. Leaf inoculation was based on a protocol originally developed at the Leibniz Institute of Plant Genetics and Crop Plant Research and adapted to our conditions. Three- to four-weeks-old wheat leaves were randomly selected and cut into 4 cm lengths. A hole was punctured in the middle of the leaf segment and inoculated with a 30 µl spore suspension. Three biological replicates consisting of each of two technical replicates were inoculated with either *F. graminearum* or *F. culmorum*. A total of 360 leaf segments were placed in water agar and incubated in phytotron for four days. The necrosis was observed in each of the 360 leaf segments and scored visually.

### Image analysis and morphotyping

Isolate pictures were analyzed using the open-source Fiji from ImageJ (**Schindelin et al., 2012**). To minimize noise from the background, the region of interest (ROI) was defined manually with the circle tool and added to the ROI manager (“Analyze/Tools/ROI manager/Add” and “Edit/Clear outside”). Once the ROI was defined, general color data was extracted (“Analyzed/color histogram”). After this first step, the image was split into three different channels (blue, green, and red), and the different features of interest were extracted for each channel (“ROI manager/Measure”). To proceed with texture analysis of each channel image, we analyzed the image with the GLCM texture macro, defining the size of the step in pixels as 1 and the direction of the step as 0 degrees (“Plugins/GLCM texture”). Finally, the image type was changed to 32-bit (“Image/Type/32-bit”) to perform the last step of the analysis. We selected *SurfCharJ1q* macro (“Plugins/ SurfCharJ1q”) to analyze roughness features. At the end of the analysis, each sample had 90 different variables to be studied.

### Statistical analysis

All statistical analyses were performed with R (version 4.2.2.) in Rstudio environment (**R Core Team, 2021**). The packages used for the analyses were: *caret, cluster, corrplot, datawizard, factoextra, faraway, readxl,* and *stats*. Data collected from ImageJ was normalized throughout the Z-scoring transformation. After data transformation, a linear model was adapted to analyze the putative correlation among the studied variables. Variables without changes among samples or with a correlation coefficient above 0.9 were removed. After this first filtering step, we performed a second selection, deleting variables with the highest correlation coefficients and variance inflation factors (VIF) values. Principal Component Analysis (PCA) was performed with the filtered and normalized data. To perform clustering analysis, agglomerative nesting (AGNES) and divisive analysis (DIANA) algorithms were applied for hierarchical clustering over our filtered data. Finally, *kmeans* and *medoids* analysis was performed with a predefined number of clusters of 2. *Kmeans* analysis was performed with *Hartigan* and *Wong* algorithm and Euclidean distance was selected for *medoids* analysis. The aggregation level of each cluster was measured throughout their silhouette value.

## Results

### Overall diversity of the wheat mycobiome library

In this study, hundreds of colonies were isolated from spikes and leaves collected over three field campaigns in eight counties across Sweden (Figure 1A). Of these, several isolates did not make it to the final set due to practical issues with growth, bacterial contaminations, failure of the genotyping at the PCR or the fragment-sequencing step. Still, we were able to recover a final set of 940 successfully genotyped samples which was the basis for the following analyzes. First, we concatenated all the genotypic data and evaluated the overall diversity of the resulting fungal collection in terms of genus diversity (Figure 1B). Here, it is important to read the following primarily in terms of qualitative diversity, considering the inherent bias due to the isolation method, and where abundance is primarily meant in terms of frequency of isolation. We were able to recover taxa from 14 different genera with a minimum of two representative (i.e. independently recovered isolates). Of these, the most abundant genera included *Fusarium* sp., the ubiquitous *Alternaria* sp., and to a lesser extent *Zymoseptoria* sp., *Pyrenophora* sp., *Microdochium* sp., and *Parastagonospora* sp., which is consistent with the fact that the sampling was mainly targeted to FHB, Leaf Blotch diseases of wheat. We therefore conclude that the resulting isolate library is relevant to the diseases covered by the survey.

We then assessed the species diversity of *Fusarium* sp. based on further analysis of ITS sequence specificity, markers, and colony morphology. Considering the limited species resolution we can obtain from ITS markers, even with all these approaches combined, frequencies must be here taken as an indication. We were able identify a several taxa typically associated with FHB in wheat, with the most common species including *F. culmorum*, *F. graminearum*, *F. poae*, and *F. avenaceum* (Figure 1C). We conclude that the collected data indicate a diversity of taxa typically implicated in head diseases of wheat present in Swedish wheat fields, representing a potential risk for disease outbreaks.

Finally, we assessed the overall diversity of the library in terms of the representation of disease-causing agents as well as potential antagonists potentially corresponding to candidate biological control agents. We reasoned that biological control agents often co-exist with their targets in the same ecological niches and may thus be co-isolated from the same tissues. To assign putative species name, we combined a series of classical pathological, symptom, and morphological criteria based on the observation of symptoms in the original material, and whenever possible, fruiting bodies, colony morphology, and spore morphology. We were able to satisfactorily identify 16 different species. Of these, 10 taxa correspond to pathogens commonly implicated in wheat diseases affecting the seeds, the leaves, and the spikes, which is consistent with the wheat organs/tissues surveyed in this study (Table 1A, 1B). We also note that head blight and leaf blotch diseases are well represented in this list which is again in line with the aim of this survey. Two putative species, *Nigrospora oryzae* and *Stemphylium vesicarium*, have been described as potentially new wheat pathogens (**Liu et al., 2021, Mohamed et al., 2022**), which we have classified as candidate emerging diseases for surveillance (Table 1B). We also found four taxa that may represent potential biological control agents which were assigned based on previous studies (Table 1C). We argue that such isolates represent an added value for the survey aiding the identification of potentially beneficial microorganisms that could be evaluated for their biological control properties.

**Table 1.** Summary of possible contributions of different taxa to the wheat mycobiome.

| Species | Predicted contribution | Reference |
| --- | --- | --- |
| **A. Common wheat pathogens** |  |  |
| *Alternaria alternata* | Black point | Somma et al. (2019) |
| *Alternaria infectoria* | Black point | Somma et al. (2019) |
| *Fusarium avenaceum* | Head blight | Karlsson et al. (2021) |
| *Fusarium culmorum* | Head blight | Karlsson et al. (2021) |
| *Fusarium graminearum* | Head blight | Karlsson et al. (2021) |
| *Fusarium poae* | Head blight | Karlsson et al. (2021) |
| *Microdochium nivale* | Fusarium Head Blight | Karlsson et al. (2021) |
| *Parastagonospora nodorum* | Septoria nodorum blotch | Bertazzoni et al. (2021) |
| *Pyrenophora tritici-repentis* | Tan spot | Moolhuijzen et al. (2022) |
| *Zymoseptoria tritici* | Septoria tritici blotch | Goodwin et al. (2011) |
| **B. Potentially emerging diseases** |  |  |
| *Nigrospora oryzae* | Seed pathogen | Eken et al. (2016) |
| *Stemphylium vesicarium* | Leaf spot | Farag et al. (2022) |
| **C. Candidate biological control agents** |  |  |
| *Aureobasidium pullulans* | Beneficial \| Antagonist | Wachowska et al. (2014) |
| *Epicoccum nigrum* | Beneficial \| Antagonist | Ogorek et al. (2011) |
| *Meyerozyma guilliermondii* | Beneficial \| Antagonist | Kthiri et al. (2021) |
| *Penicillium chrysogenum* | Beneficial \| Antagonist | Barna et al. (2008) |

### Inventory of pathogenic and endophytic isolates associated with Leaf Blotch in Sweden

For the leaf mycobiome associated with Leaf Blotch diseases, we recovered isolates from typical blotch symptoms as well as from symptomless tissues. We deduced that symptomless tissues may be the habitat for latent pathogens, endophytic fungi that could be antagonistic to the pathogenic, or harbor pathogens surviving as silent endophytes.

Overall, *Alternaria* species dominated the infection, which is consistent with the fact that they are known to be ubiquitously found as weak pathogens or saprophytes on different plant tissue (Figure 2A). *Alternaria* sp. apart, the most frequently isolated species were *Zymoseptoria* sp., *Pyrenophora* sp., and *Parastagonospora* sp. Evidence from the symptoms where these taxa were isolated, combined with colony morphology, and the observation of typical fruiting bodies indicated that the species are *Zymoseptoria tritici*, *Pyrenophora tritici-repentis*, and *Parastagonospora nodorum*, respectively, all of which being prominent causal agents of blotch diseases in wheat. We here conclude that the collection isolated from leave does represent the diseases targeted by the survey. We then differentiated the isolates recovered from typical blotch symptoms (Figure 2B), from those recovered from symptomless tissues (Figure 2C). *Alternaria* sp. apart, *Zymoseptoria* sp., *Pyrenophora* sp., and *Parastagonospora* sp were the most frequently isolated species from blotch symptoms, with further symptomatic and morphological inspection indicated these isolates correspond to the species *Zymoseptoria tritici*, *Pyrenophora tritici-repentis*, and *Parastagonospora nodorum*, respectively.

**Figure 2.**
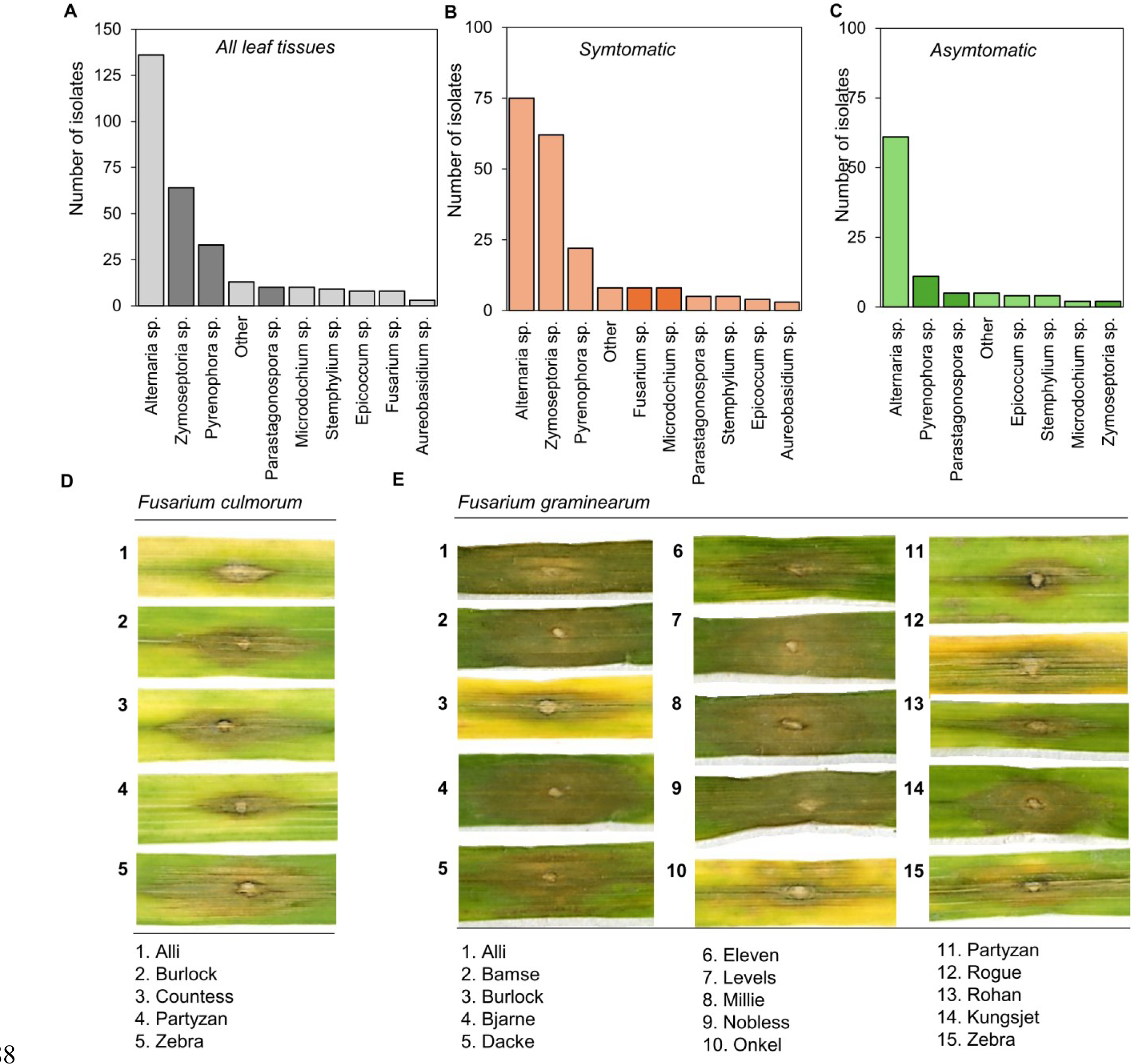
Summary of the wheat leaf mycobiome. (A) Overall diversity of the fungal taxa isolated from leaf tissues. (B) Summary of the fungal taxa recovered from tissues with typical leaf blotch symptoms. (C) Summary of the fungal taxa recovered from asymptomatic tissues from leaf blotch infected material. (D) Point leaf inoculations of different spring wheat commercial cultivars with *Fusarium culmorum* (see Methods). (E) Point leaf inoculations of different spring wheat commercial cultivars with *Fusarium graminearum* (see Methods).

Interestingly, we have also isolated taxa typically implicated in FHB from the genera *Fusarium* sp. and *Microdochium* sp.. Of these, several isolates had typical morphological attributes of *F. culmorum*, *F. graminearum*, and *Microdochium nivale*. In comparison, the most frequently isolated taxa from asymptomatic tissues apart from *Alternaria* sp. were *Pyrenophora* sp., and *Parastagonospora* sp., but also *Zymoseptoria* sp. and *Microdochium* sp. This data suggest that members of the FHB complex can possibly survive as latent pathogens or silent endophytes in the leaves and potentially cause blotch-like symptoms that can be confound with the typical blotch pathogens *Z. tritici*, *P. tritici-repentis*, and *P. nodorum*.

We then tested the capacity of *F. culmorum*, *F. graminearum* to cause blotch-like symptoms on wheat leaves. To do so, we performed point inoculation of leaf segments harvested at the seedling stage and visually assessed the development of necrotic lesions (see Methods). We inoculated 30 commercial spring wheat cultivars commonly grown in Sweden (Figure 2D, 2E). We found that *F. culmorum* caused necrotic lesions on at least 5 out of the 30 commercial cultivars tested (Figure 2D), while *F. graminearum* caused necrotic lesions on 15 commercial cultivars (Figure 2E). These results substantiate the hypothesis that *F. graminearum* and *F. culmorum* can potentially cause leaf blotch-like symptoms in Swedish fields on several of the commonly grown spring wheat varieties.

### Inventory of pathogenic and endophytic isolates associated with FHB in Sweden

For the spike mycobiome, we recovered isolates from seeds as well as from glumes and searched for both pathogenic and non-pathogenic taxa. Our reasoning was that glumes may offer a secondary habitat for wheat ear pathogens or potential antagonists competing for the same ecological niche in the seed vicinity. In contrast to the leaf mycobiome, *Fusarium* species dominated over *Alternaria* sp. (Figure 3A). We then investigated the differences in the qualitative diversity between the taxa recovered from seeds vs. glumes (Figure 3B, 3C). Interestingly, a much larger diversity of taxa was isolated from seeds, while Fusarium sp. dominated in glumes as compared where a limited diversity of taxa was isolated (Figure 3B, 3C). We also observed that *Fusarium* species fully occupy the seed habitat, with the glumes possibly acting as a secondary niche where *Fusarium* sp. specifically seem to outcompete other fungal taxa, such as *Alternaria* sp which has been isolated ubiquitously from all tissues analyzed in this study except glumes.

**Figure 3.**
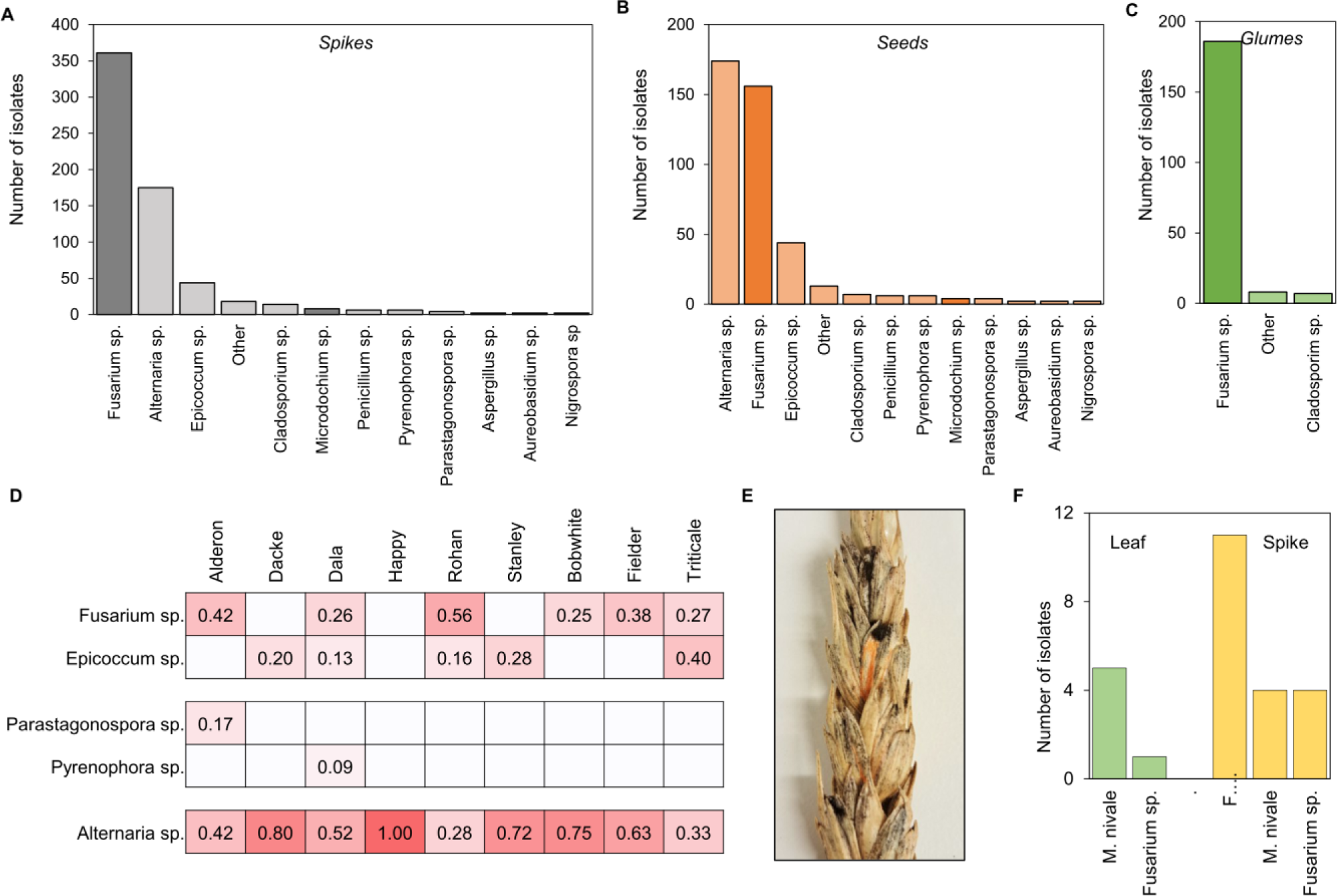
Summary of the wheat head mycobiome. (A) Overall diversity of the fungal taxa isolated from spikes. (B) Summary of the fungal taxa recovered from seeds. (C) Summary of the fungal taxa recovered from glume tissues. (D) Analysis of possible association between fungal taxa and wheat cultivars. (E) Photograph of typical Fusarium Head Blight symptoms on a wheat spike collected from the most northern location in our survey: Piteå. (F) Summary of the fungal taxa isolated from typical leaf blotch and head blight symptoms from samples collected in Piteå.

We then expanded this analysis to possible differences between cultivars. We used a subset of samples recovered from a small propagation plot, where isolations were exclusively carried out with seeds. First, we concatenated the data by cultivar and genus with at least two representatives (Figure 3D), Additionally, we also included isolates from *Epicoccum* sp. which we could isolate from leaves, but also and mostly from seeds. Our reasoning was that *Epicoccum* sp. had been previously described as being associated with FHB, however, while a role as a possible antagonist/biocontrol agent has been suggested (**Karlsson et al., 2021**), its contribution to the FHB complex is still poorly understood. We found that *Alternaria* sp. could be isolated from most, if not all, cultivars, which is consistent with their characteristic as ubiquitous plant-associated fungi, sometimes acting as weak pathogens. However, the distribution of *Fusarium* sp. and *Epicoccum* sp. was more scattered across cultivars (Figure 3D).

We finally investigated the specific case of FHB-infected ears (Figure 3E) and leaves with typical blotch symptoms that were collected from the most northern location in our study: Piteå (Norrbotten county, Figure 1A). Interestingly, we mostly recovered *Michrodochium* sp. colonies (with typical features of *M. nivale*) from the leaf blotch symptoms and *F. avenaceum* and *Michrodochium* sp. from the FHB-infected ears (Figure 3G). These results suggest that in these northern latitudes, blotch symptoms can be probably caused by *Michrodochium* sp., which is the common agent of the Snow Mold disease of wheat, while FHB is possibly caused by the weaker pathogens *F. avenaceum* and *M. nivale*. We conclude that such observations demonstrate the usefulness of such surveys to shed light on possible changes in the disease landscape and alert on the possible re-emergence of diseases.

### Morphotyping of *Fusarium* sp. and *Epicoccum* sp. isolates

In a last series of analyses, we took advantage of the library to explore the possibilities of morphotyping to rapidly sort and identify fungal species, with the prominent addition of modern image analysis technologies. We inferred that the capacity to rapidly sort and classify fungal taxa as soon as they are available as colonies growing on a plate will be a handy tool to accelerate and scale up the diagnostics, which would also allow us to target specific taxa or specific morphotypes. To do so, we aimed to provide a proof-of-concept using *Fusarium* sp. and *Epicoccum* sp. as targe taxa to differentiate solely using imaging.

A pipeline was designed to extract different image attributes from the isolate pictures using advanced functions in the imageJ image analysis suite (see Methods). A Principal Component Analysis (PCA) showed that although the subset of selected image variables still shows some level of correlation (Figure 4A), they can still explain 56.7 % of the observed variability (Figure 4B). The image features with the highest contribution to the ‘*Dimension 1*’ is the mean blue channel, and the blue and green intensity density (Figure 4C), meanwhile the highest contributors to ‘Dimension 2’ are the minimum green intensity and the FAD values for red, blue, and green (Figure 4D). These results suggest that the selected characteristics could represent a balanced mathematical description of the distribution of our samples. However, we cannot exclude further optimization considering the inherent correlation between some visual features (such as color) and the fact that some image features contribute more to the total variability. Finally, with this still basic analysis, we could delineate two clear but slightly overlapped clusters (Figure 4B) differentiating *Fusarium* sp. from *Epiccocum* sp. isolates. In order to improve the clustering of our data, we applied Agglomerative Nesting (AGNES) (Figure 4E) and Divisive Analysis Clustering (DIANA) (Figure 4F) algorithms (see Methods). The AGNES algorithm resulted in a dendrogram with an agglomerative coefficient of 0.78; meanwhile, the DIANA algorithm gave a higher value of 0.82, suggesting a solid clustering structure in our sample distribution. Although the clustering does not differentiate between different *Fusarium* sp. species, the distribution shows that we can reasonably differentiate between the genus *Epicoccum* sp. and *Fusarium* sp. with only six miscalls.

**Figure 4.**
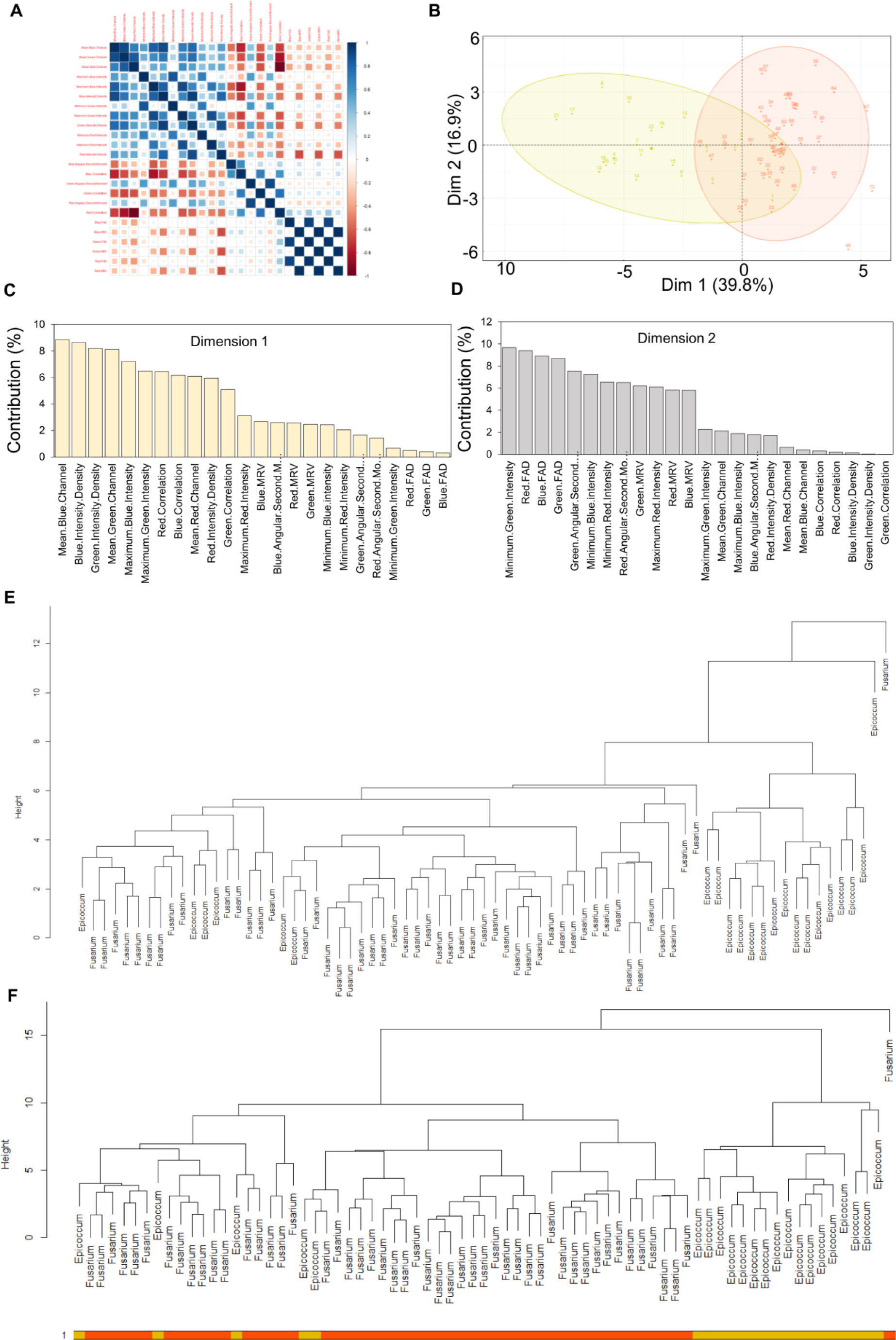
Feature extraction for morphotyping and dendrogram classification. (A) Correlation matrix of selected variables for image analysis of *Epicoccum* sp. and *Fusarium* sp. blue color (values close to 1) indicates positive correlation meanwhile red color indicates negative correlation. (B) Principal component analysis plot of visual data extracted from *Epicoccum* sp. and *Fusarium* sp. Isolate images. The color yellow represents *Epicoccum* sp. isolates, and the red color represents *Fusarium* sp. isolates. Circles represent putative clusters. Dimension 1 and dimension 2 separate isolates based in the values of 24 different visual characteristics. (C) Weights of the main explanatory variables for the first PCA dimension. (D) Weights of the main explanatory variables for the second PCA dimension. (E) Dendrogram clustered by AGNES algorithm built with Unweighted Pair Group Method with Arithmetic Mean (UPGMA). (F) Dendrogram clustered by the DIANA algorithm. The yellow color represents *Epicoccum* sp. isolates, and the red color indicates *Fusarium* sp. isolates.

Considering the quality of the data, we aimed at further clustering using *K-means* (Figure 5A) and *Partition Around Medoids* (*PAM*) algorithms (Figure 5B), and calculated the average silhouette value for each cluster created (Figure 5C, 5D). Although the global accuracy increased was improved to 0.918 (*K-means*) and 0.904 (*PAM*), we still had 6 miscalls with the *K-means* and 7 with the *PAM*, which is consistent with the average silhouette value of 0.32 for the *K-means* clusters (Figure 5C) and 0.3 for the *PAM* clusters (Figure 5D). Interestingly, it seems that algorithms find it harder to differentiate some *Epicoccum* sp. isolates from *Fusarium* sp. (*K-means*: 0.739, PAM: 0.739) while the sensitivity for *Fusarium* (i.e. the capacity to properly identify *Fusarium* sp.) is higher in comparison to *Epiccocum* (*K-means*: 0.980, PAM: 0.961. Therefore, despite the small dataset used in this analysis, we could still provide proof of principle that imaging is a promising tool for advanced morphotyping and the identification of specific fungi relevant to wheat.

**Figure 5.**
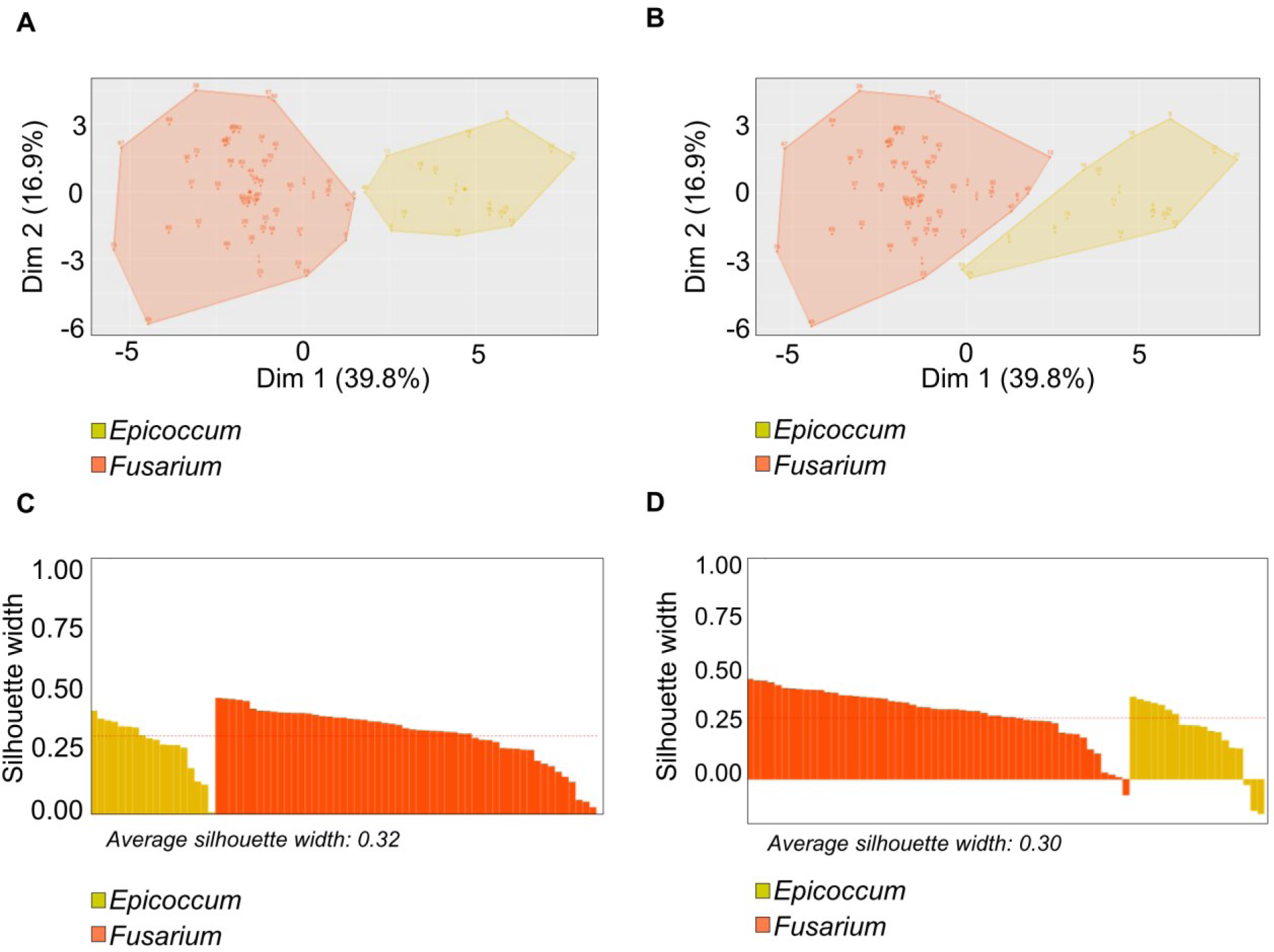
K-mean and PAM clustering of *Epicoccum* sp. and *Fusarium* sp. Isolates. (A) Cluster analysis performed by K-means algorithm. The pre-selected number of clusters was defined as 2. Yellow represents the putative cluster for *Epicoccum* sp. isolates and red color represents the putative cluster for *Fusarium* sp. (B) Cluster analysis performed by PAM algorithm. The number of clusters was predefined as 2. Yellow area represents *Epicoccum* sp. isolates and the red area shows *Fusarium* sp. (C) Silhouette values for each isolate and average silhouette from cluster analysis with the K-means algorithm. (D) Silhouette values for each isolate and average silhouette from cluster analysis with the PAM algorithm. In (C) and (D) samples aggregated as *Epicoccum* sp. are represented with yellow bars meanwhile samples aggregated as *Fusarium* sp. are represented as red bars.

## Discussion

### Wheat pathogens occupy a variety of tissues and niches

In this study, we recovered typical wheat pathogens implicated in head blight and leaf blotch diseases from a variety of tissues, including seeds, glumes, symptomatic leaf tissues, and asymptomatic leaves. We had cases where *Fusarium* taxa typically implicated in FHB were isolated from blotch symptoms in the leaves and other cases where typical blotch pathogens such as *P. tritici-repentis*, and *P. nodorum* were isolated from wheat ears. Furthermore, typical leaf pathogens were also isolated as latent pathogens or silent endophytes from asymptomatic tissues. Fusarium species are well known as ubiquitous plant colonizers that can survive in a variety of plant tissues, and also in the soil (**Karlsson et al., 2021**). In disease complexes such as FHB, where 17 different *Fusarium* and non-*Fusarium* taxa can be implicated, the role of such secondary niches is poorly understood (**Karlsson et al., 2021**). From a population perspective, the capacity of pathogens to survive in tissues or organs that are not their primary habitat may increase the fitness of the pathogen population in the field by maintaining living inoculum, throughout the field seasons and particularly when conditions are unfavorable. This was, for instance, observed in the case of *Z. tritici*, where it was shown that avirulent isolates can still survive and mate on resistant wheat hosts as symptomless endophytes or epiphytes (**Fones et al., 2023**), thus contributing to pathogen fitness and adaptation through the maintenance of genetic diversity in the field population. It was also shown that *Z. tritici* can induce systemic responses leading to whole plant metabolic changes leading to increased susceptibility to secondary infection with a non-adapted pathogen (**Seybold et al., 2020**). This is major because such mechanisms could also occur among different isolates of the same pathogen population or different species of a disease complex, thus providing a possible explanation for the functional relevance of secondary niches and the ubiquitous nature of the taxa implicated in FHB.

### The wheat mycobiome as a source for candidate biological control agents

Microbial biological control agents are considered an important tool to reduce chemical inputs in crop disease management. A variety of microbial agents with antagonistic properties against plant pathogens has been described, and variety of mode of action underlying such effect has been reported including antibiosis, parasitism, niche competition, or resistance priming in the host (**Köhl et al., 2019**). Microbial biological control agents that correspond to naturally occurring competitors are commonly found in the same ecological niches as their target pests. In this study, a few candidate biological control agents have been identified (Table 1C). Of these, *Epicoccum* sp. have been shown to be a promising biological control agent against wheat stripe rust as well as *F. graminearum*, while *Aureobasidium pullulans* and *Meyerozyma guilliermondii* have been described as biological control agents against post-harvest Fusarium crown rot diseases, respectively (**El-Sharkawy et al, 2023; Wachowska et al., 2016; Kthiri et al., 2021**). These results further demonstrate the usefulness of such surveys in the broader context of IPM, by facilitating the identification of relevant candidate biological control agents better adapted to the Swedish conditions.

### Wheat disease diagnostics and its importance for management

The beneficial vs. pathogenic lifestyle of a plant colonizing fungus can be determined mainly by the host and the ecological niches it occupies. For instance, the biological control fungus *Clonostachys rosea*, a naturally occurring root endophyte, has been shown to be effective against several foliar plant pathogens. However, *C. rosea* has also been reported as the causal agent of root rot, crown rot, and wilt-type diseases in garlic, faba bean, avocado, strawberry, sugar beet, and medicinal plants such as *Angelica sinensis* and *Astragalus mongholicus* (**Ma et al., 2020; Qi et al., 2022; Diaz et al., 2022; Afshari et al., 2017; Coyotl-Pérez et al., 2007; Zhang et al., 2022; Haque and Parvin, 2020, Farhaoui et al., 2023**). Similarly, *Epicoccum* sp. have been described as a potent biological control agent against wheat stripe rust. However, *E. nigrum* is also the causal agent of the brown spot disease in *Solanaceae* (**Jin et al., 2023**). These reports highlight the importance of continuous monitoring of field populations and careful, systematic identification of the causal agents for identifying potential lifestyle shifts in microbial populations.

Shifts from a beneficial endophytic lifestyle to pathogenicity through host switch, niche displacement, and changes in the microbiome have been well documented for several prominent diseases such as *Diplodia sapinea,* the causal agent of diplodia tip blight in Scots pine, and *Sarocladium oryzae* the causal agent of rice sheet rot (**Blumenstein et al., 2022; Peeters et al., 2021**). Therefore, the designation of beneficial endophyte/candidate biological control agents must be taken within the context of the actual contribution of the species to the wheat mycobiome, meaning that both disease-suppressing antagonistic and disease-promoting synergistic properties must be tested. This is directly relevant to taxa such as *Epicoccum* sp. and *Alternaria* sp., considering their frequent association with FHB and our poor understanding of how such mycobiome is assembled (**Karlsson et al., 2021; Schiro et al., 2018; Nazabanita et al., 2022; Rojas et al., 2020**).

In this context, changes in the composition of the disease complex are also relevant, with the example of FHB from Piteå, which seems to be caused by *F. avenaceum* and *M. nivale* (Figure 3F, 3G) and not by *F. graminearum* and *F. culorum* as in the majority of the FHB tested samples (Figure 3A, 3B, 3C). In contrast, we also recovered those typical FHB causal agents, *F. graminearum* and *F. culorum*, from typical necrotic and chlorotic symptoms reminiscent of septoria leaf blotch diseases. These results highlight the complexity of the wheat mycobiome and the limits of visual scoring considering the complexity and often the convergence of disease symptoms caused by yet phylogenetically distant taxa. We argue that such observations further highlight the importance of field surveys as a tool for continuous identification and re-identification of the actual disease-causing agents present in the field.

### A basis for targeted monitoring of potentially emerging wheat diseases

The emergence of new diseases leading to global epidemics is relatively common in wheat, with stem rust and wheat blast being two prominent examples. Wheat has also been the host for the emergence of two new pathogens that did not existence before namely triticale powdery mildew and wheat blast. Arguably, the large reservoir of wheat pathogens present in the agro-ecosystem is a challenge requiring continuous monitoring for early detection of potentially new, emerging diseases. In this context, two taxa identified in our survey are of particular interest: *N. oryzae*, and *S. vesicarium* (Table 1B). *N. oryzae* is an ascomycete fungal pathogen and the causal agent of panicle branch rot disease on rice (*Oryza sativa*) (**Liu et al., 2021**) In 2016, *N. oryzae* was reported to cause black lesions on the hypocotyls of wheat seedlings in Kazakhstan, suggesting that some isolates of *N. oryzae* (a rice pathogen) may have adapted to be pathogens on wheat, which is reminiscent of the emergence of wheat blast (**Eken et al., 2016**). Similarly, *S. vesicarium* has been reported to cause leaf spots on wheat in Egypt (**Mohamed et al., 2022**), and to our knowledge, these pathogens have never been described in Sweden before. We conclude that such results further highlight the relevance of this work since we can now provide a rational basis for risk assessment and targeted disease monitoring of critical pathogens in Swedish wheat fields.

## Author Contributions

SB and HV led the research. SB, HZ, and JZ designed the research and developed the library. HK, EV, MACG, HV, SB did laboratory experiments. HK, MACG, SB, DD, and FD developed infection assays. MACG and SB developed the morphotyping. HK, EV, HV, and SB analyzed the genotyping data. MACG performed statistical analyzes. MACG and SB edited the figures.

## Acknowledgments

We would like to acknowledge the support of the Plant Protection Extensions ‘Växtskyddscentralerna’ of the Swedish Board of Agriculture ‘Jordbruksverket’ (https://jordbruksverket.se/) with field sampling across Sweden. We would like to thank Anders Lindgren, Lina Norrlund, and Elizabeth Lovetta for supporting and coordinating sample collection and shipment. We thank Eula Gems Oreiro, Björn Andersson from the Department of Forest Mycology and Plant Pathology (SLU) for their help handling blotch infected samples and their contribution to maintaining the library. We also thank Fredric Hedlund and Ayano Tanaka for their technical assistance with the growth facilities and the SLU BioCentrum. We finally thank Katarina Ihrmark and Maria Jonsson for their technical assistance.

